# ACh signaling modulates activity of the GABAergic signaling network in the basolateral amygdala and behavior in stress-relevant paradigms

**DOI:** 10.1101/2022.02.08.479551

**Authors:** Yann S. Mineur, Tenna N. Mose, Kathrine Lefoli Maibom, Steven T. Pittenger, Alexa R. Soares, Hao Wu, Yaqing Huang, Marina R Picciotto

**Affiliations:** Department of Psychiatry, Yale University School of Medicine, 34 Park Street, 3^rd^ Floor Research, New Haven, CT 06508, USA; Interdepartmental Neuroscience Program, Yale University School of Medicine, 34 Park Street, 3^rd^ Floor Research, New Haven, CT 06508, USA; Experimental Pathology Graduate Program, Yale University School of Medicine, 34 Park Street, 3^rd^ Floor Research, New Haven, CT 06508, USA

## Abstract

Balance between excitatory and inhibitory (E/I) signaling is important for maintaining homeostatic function in the brain. Indeed, dysregulation of inhibitory GABA interneurons in the amygdala has been implicated in human mood disorders. We hypothesized that acetylcholine (ACh) signaling in the basolateral amygdala (BLA) might alter E/I balance resulting in changes in stress-sensitive behaviors. We therefore measured ACh release as well as activity of calmodulin-dependent protein kinase II (CAMKII)-, parvalbumin (PV)-, somatostatin (SOM)- and vasoactive intestinal protein (VIP)-expressing neurons in the BLA of awake, behaving mice. ACh levels and activity of both excitatory and inhibitory BLA neurons increased when animals were actively coping, and decreased during passive coping, in the light-dark box, tail suspension and social defeat. Changes in neuronal activity preceded behavioral state transitions, suggesting that BLA activity may drive the shift in coping strategy. In contrast to exposure to escapable stressors, prolonging ACh signaling with a cholinesterase antagonist changed the balance of activity among BLA cell types, significantly increasing activity of VIP neurons and decreasing activity of SOM cells, with little effect on CaMKII or PV neurons. Knockdown of α7 or β2-containing nAChR subtypes in PV and SOM, but not CaMKII or VIP, BLA neurons altered behavioral responses to stressors, suggesting that ACh signaling through nAChRs on GABA neuron subtypes contributes to stress-induced changes in behavior. These studies show that ACh modulates the GABAergic signaling network in the BLA, shifting the balance between SOM, PV, VIP and CaMKII neurons, which are normally activated coordinately during active coping in response to stress. Thus, prolonging ACh signaling, as occurs in response to chronic stress, may contribute to maladaptive behaviors by shifting the balance of inhibitory signaling in the BLA.

## INTRODUCTION

Exposure to stressors and traumatic life experiences contributes significantly to the onset of mood disorders ^1^. The amygdala plays a central role in emotional processing and is one of the primary brain regions associated with stress responses ^2, 3^. Hyperactivity of the amygdala is observed in individuals with mood disorders and may contribute to the pathophysiology of stress-related disorders, including depression and anxiety ^4^. This is in line with the idea that depression is a disease of disrupted neurocircuitry ^5^. Depression and anxiety disorders may therefore be characterized as maladaptive behavioral responses to stress.

Pre-clinical studies corroborate human studies, pinpointing the amygdala as an important brain area mediating stress responses, as well as a region involved in mediating anxiolytic- and antidepressant-like effects of cholinergic antagonists ^6^. In particular, the basolateral amygdala (BLA) receives substantial cholinergic input and is involved in aversive learning and anxiety-like behaviors ^7, 8^. During exposure to restraint stress, acetylcholine (ACh) levels increase in multiple regions of the limbic system, including the BLA ^9^. Previous studies have demonstrated that administration of the non-selective nicotinic acetylcholine receptor (nAChR) antagonist mecamylamine, as well as genetic knockdown (KD) of β2-containing or α7 nAChR subunits in the BLA, can reduce anxiety- and depression-like behaviors in mice ^10^ and decrease BLA neuronal activity ^10, 11^. These data suggest that decreasing cholinergic signaling through nAChRs in the BLA can provide resilience to stress-induced behaviors. However, ACh can also improve cognition and the valence of positive stimuli ^7, 12, 13^, indicating that optimal homeostatic levels of ACh are required for proper modulation of attention, learning, and stress responses ^14^.

Cholinergic regulation of network excitability in the BLA can result from direct ACh input to principal neurons ^15^ or via modulation of GABAergic interneurons that form local inhibitory circuits ^16^. Thus, behavioral responses to stressful events depend on how cholinergic transmission alters the individual neuronal cell types in the network. The BLA consists of excitatory glutamatergic principal projection neurons (PN) that express calcium- and calmodulin-dependent protein kinase II (CaMKII) and a complex network of inhibitory GABA interneurons. GABAergic subtypes include parvalbumin (PV)-expressing cells that form connections onto cell bodies of PNs and onto somatostatin (SOM)-expressing inhibitory interneurons ^17^. SOM interneurons, in turn, target both PV and vasoactive intestinal peptide (VIP)-expressing GABA interneurons. Finally, VIP interneurons target PNs, and innervate PV, SOM, as well as other VIP interneurons ^17–19^. The interconnections between cell types in the local GABAergic circuitry of the BLA provides the possibility for neuromodulatory inputs, such as ACh, to fine-tune network excitability, which is critical for proper behavioral responses to environmental stressors.

We therefore measured the activity of each of these neuronal subtypes (CaMKII, PV, SOM, VIP) in freely moving mice during stress-induced behaviors and evaluated the effect of prolonging ACh signaling on the balance of activity in the network. In order to determine the functional consequence of ACh inputs to BLA on the network, we genetically downregulated expression of the primary nAChR subtypes in the BLA (β2 and α7 nAChRs) in CaMKII or GABA neurons, and then selectively in PV, SOM or VIP neuronal subtypes, to determine whether cholinergic transmission differentially alters stress-relevant behaviors through different neuronal subtypes in the BLA network.

## METHODS

### Animals

GAD-, PV-, SOM-, VIP-, and CaMKII-Cre mice were backcrossed in our facility onto the C57BL/6J background and adult, male mice were used for all experiments.

GAD1(GAD65)-Cre mice were originally obtained from Dr. Lilliana Minichiello ^20^. CaMKIIα-Cre mice were originally obtained from Dr. Gunther Schütz ^21^. PV-, SOM- and VIP-Cre mice were obtained from Dr. Nobuaki Tamamaki ^22^. Male C57BL/6J mice were purchased from the Jackson Laboratories (Bar Harbor, ME, USA). Animals were housed in a vivarium at ~21°C on a 12-hour light-dark cycle from 7 am to 7 pm. Food and water were available *ad libitum*. Animals were group-housed unless separated for humane concerns. Animals from each strain were randomized into the different experimental groups.

Retired male CD1 breeders purchased from Charles River Laboratories (Wilmington, MA, USA) were used in the Social Defeat (SD) paradigm as aggressors. CD1 mice were single-housed under the same conditions described above. All procedures involving animals were approved by the Yale University Committee on the Care and Use of Animals and were carried out in accordance with the Guidelines in the National Institutes of Health Guide for the Care and Use of Laboratory Animals.

### Stereotaxic surgery

Animals were maintained under isoflurane anesthesia (Zoetis, Kalamazoo, USA) in a stereotaxic frame (Kopf Instruments). Anesthesia was monitored throughout the surgical procedure and reflexes were checked frequently by toe pinch. An incision was made to expose the skull and one (for fiber photometry) or two (for bilateral viral knockdown) small holes were drilled into the skull to allow access to the brain. A 26-gauge blunt-tipped Hamilton syringe was then used to deliver viruses carrying the appropriate constructs into the BLA (coordinates: −1.8 mm AP, ±3.4 mm ML, 5.2 mm DV from bregma) at a rate of ~0.1 μL/min. To minimize backflow, the needle was kept in place for an additional 5 min after injection and lowered 0.1 mm just before removal.

Animals recovered in a cage on a heating pad before being returned to their home cage. Animals were monitored post-surgery and were treated with 5 mg/kg carprofen (Butler Schein Animal Health, Columbus, OH, USA) for 46 hr. Carprofen was diluted in phosphate-buffered saline (PBS) (pH 7.4) for the injection solution and injected intraperitoneally (i.p).

### Fiber photometry (**Fig. 1A**)

0.3 μL of pAAV.Syn.Flex.GCaMP6f.WPRE.SV40 (Addgene 100833-AAV1), pENN.AAV.CamKII.GCaMP6f.WPRE.SV40 (Addgene 100834-AAV1), or AAV1-GRAB_ACh_ 3.0 ^23^ was injected into the right BLA. An optical fiber (400 μm diameter/6 mm length; Doric Sciences, Quebec, Canada) was implanted 0.1 mm above the injection site and maintained in place with adhesive dental cement (C&B-Metabond, Parkell, New York U.S.A). Cement was allowed to harden before the incision was closed and the animal was allowed to recover for at least 3 wk before testing to ensure proper healing and adequate GCaMP6f or GRAB_ACH_ expression (**Fig. 1B**).

**Figure 1:**
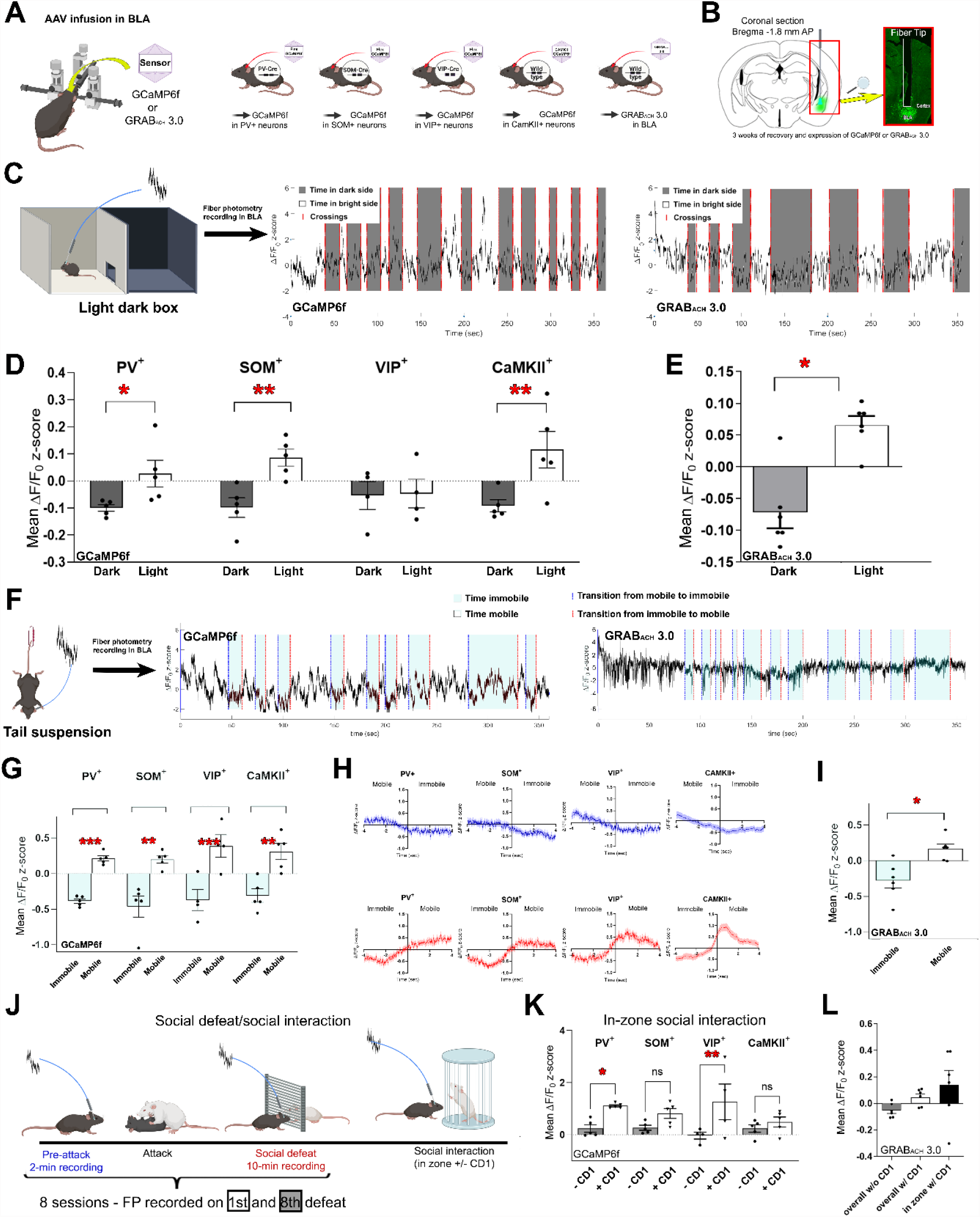
BLA infusion of Cre-dependent (FLEX) GCaMP6f in Cre-expressing mice, CAMKII-dependent GCamp6f and GRAB_ACh_ 3.0 in wild-type mice (**A**). Fiber placements and viral expression (GFP) were verified for all animals following experimentation, and animals were tested in a battery of behavioral assays staring ~3 weeks post-surgery to allow for efficient expression of GCaMP6f or GRAB_ACh_ 3.0 fluorescent sensors (**B**). Fiber photometry (FP) recording in the light-dark box (**C**) and example traces for GCaMP6f and GRAB_ACh_ 3.0 fluorescence measurements. FP data were split between time in the light and dark compartments, and average z-score was then calculated for each condition. (**D**) Mean GCaMP6f z-scored ΔF/F0 for activity of each neuronal subtype when animals were in the light or dark side of the light-dark box. (**E**) Mean GRAB_ACh_ 3.0 z-scored ΔF/F0 in the BLA when animals were in the light or dark side of the light-dark box. Error bars represent SEM and solid dots are individual averages per animal. FP recording in the tail suspension test (**F**) and example traces for GCaMP6f and GRAB_ACh_ 3.0 fluorescence measurements. FP data were split between mobile and immobile states, and average z-scores were then calculated for each condition (**G**). Error bars represent SEM and solid dots are individual averages per animal. (**H**) Mean GCaMP6f z-scored ΔF/F0 across animals, recorded 4 sec prior to 4 sec after a change in mobility state in the tail suspension test. Solid lines are means across events and animals. Shaded areas are SEM. (**I**) Mean GRAB_ACh_ 3.0 z-scored ΔF/F0 in the BLA during mobile or immobile episodes in the tail suspension test. Error bars represent SEM and solid dots are individual averages per animal. N = 5 per group for each test with GcAMP6f and n = 6 for GRAB 3.0. * p < 0.05; ** p < 0.01; *** p < 0.01 Partially created with Created with BioRender.com.

### Cell type-specific nAChR knockdown

Small hairpin RNAs (shRNAs) targeting specific nAChR subtypes (AAV2-dsRed-Sico-α7-shRNA or AAV2-dsRed-Sico-β2-shRNA) were cloned into the pSICO backbone as previously described ^24–27^ and were used to knock down expression of nAChR subtypes in a cell type-selective manner. 0.5 μL of AAV2-dsRed-Sico-α7-shRNA or AAV2-dsRed-Sico-β2-shRNA was injected bilaterally into the BLA of CaMKII-, SOM-, PV- or VIP-Cre mice. A scrambled version of the shRNA was used as a control. To allow time for robust knockdown of α7 or β2* nAChR mRNA and degradation of remaining protein, mice were allowed to recover for at least eight weeks after stereotaxic surgery and before behavioral testing.

### Drug treatment

Physostigmine (Sigma) was injected i.p. at 10 mg/kg (10 mL/kg), 60 min before behavioral assays or immediately before fiber photometry recording. Before physostigmine injection, fluorescence was recorded for 10 min (Baseline), and was then recorded for 5 min following Saline injection. Results are reported as Z-scored ΔF/F calculated over the entire session (see <u>Fiber Photometry Data Processing</u> section below for more information).

### Behavioral assays

A battery of tests was performed in sequence as described below unless incompatible with the type of experimental design. 24 to 48 hr was allowed between each assay.

#### Light Dark (LD) test

The light-dark box consisted of two compartments: a dark compartment (safe) and a light compartment with a bright light bulb above (exposed). Each test subject was placed in the corner of the light compartment facing away from the opening to the dark compartment. The animal was allowed to move freely around and between the two-chambers. Following the first crossing into the dark compartment, the time spent in each compartment was measured for 6 min. Each animal was returned to a holding cage after the test until all cage mates were tested.

To accommodate the fiber in fiber photometry experiments, a small groove was made in the lid and the separating wall between the chambers, and a 3D-printed funnel allowed transitions between the compartment for animals with a connected fiber.

#### Tail suspension test (TST)

A paperclip was taped to the last 5 mm of the tail of each test animal. The mice were suspended by the tail for 6 min and time spent immobile was recorded. Immobility was defined as no movement except breathing and slight whisker twitches. Animals were returned to their home-cage after the test.

#### Forced swim test (FST)

Clear 4-liter glass beakers were filled with 2.5 L of water at room temp. Each animal was carefully placed in the water allowing them to keep their nose above water. On the first day of testing, the animal was placed in the water for 15 min. On the second day, the animal was placed in the water for 5 min ^10, 28^. The time spent immobile during the 5-min test on the 2^nd^ day was scored and used for statistical analyses. Immobility was defined as floating during which the animal made a minimal amount of movement to keep above water. Because of the of the attached fiber, the FST was not compatible with FP recording. Evaluation of immobility in the FST was therefore only carried out for knockdown experiments.

#### Social defeat

Male CD1 mice were single-housed with no cage change for 2 wk prior to use as an aggressor to allow the animal to establish its territory. Each animal was screened for aggressive behavior prior to use as an aggressor, as well as before every session of social defeat, to ensure a high level of aggression.

On the morning of the initial defeat experience, each mouse was placed into the home cage of an aggressor CD1 mouse. Mice were separated after the first attack with a metal grid leaving two thirds of the cage for the CD1 mouse. The test subject was left in the aggressor’s cage for 10 min before being returned to its home cage. The test was repeated in the afternoon and this process was continued for 4 days ^10^. Social interaction (SI) was measured on the 5th day. The SI test area consisted of an open field with a small holding square for the CD1 mouse. The test mouse was placed in the corner of the open field and allowed to explore freely for 2.5 min. A CD1 mouse was then placed in the holding square and 2.5 additional min of exploration/interaction were assessed. SI was expressed as the ratio between time spent in the 5 cm ratio area around the holding square with or without the CD1 mouse.

For FP experiments, the social defeat paradigm was performed as described above with modifications to accommodate the connected fiber. There was a “pre-attack” FP recording before the 1st and 8th defeat session in which the CD1 home cage was separated with a grid to prevent physical attack that might damage the recording fiber. The test mouse was connected to the patch cord and put inside the cage, isolated from the CD1, and fluorescence was recorded for 2 min. The test animal was then disconnected from the patch cord and put back into the CD1 home cage without the physical barrier. After the first attack, the animals were separated by a mesh grid. The test mouse was then reconnected to the patch cord, put back into the CD1 home cage and fluorescence was recorded for an additional 10 min. The 2nd to 7th defeats occurred without connection to the patch cord and consisted of only the initial attack and subsequent social defeat exposure ^29^.

#### Locomotor Activity in an Open Field

A locomotor activity assay was performed to identify any strain differences between Cre-expressing lines as well as any effects of connection of the patch cord in FP experiments. Each animal was placed in the center of a clear, clean, novel cage with no bedding (30×15×13cm3) and was allowed to move freely for 20 min with the patch cord connected. Distance moved was analyzed using Ethovision XT 10 and expressed as total distance moved in cm ^10^.

#### Open Field with pharmacological injection and FP recording

For measurements of the effect of physostigmine on BLA neuronal activity using FP, each animal was placed in the center of a clear, clean, novel cage (30 × 15 × 13 cm^3^) and allowed to move around freely for 10 min. Within ~3 min, the baseline Z-score became stable for all cell type measurements. Animals were then injected with 0.15 mg/kg Phosphate Buffered Saline (PBS) and placed back in the cage. 5 min later, animals were administered 0.15 mg/kg physostigmine, i.p. (Sigma Pharmaceuticals, North Liberty, IA, USA). After ~8 min, the onset of physostigmine-induced behavioral effects were observed. Mice displayed flattening and immobility that are typically observed after peripheral administration of anticholinesterase drugs ^28^. By ~ 30 min behavioral signs resolved, and animals began to explore the cage and move normally. FP recording continued for 45 min after the last injection.

### Confirmation of fiber implantation and viral expression/tissue processing

Animals were anesthetized with an i.p. injection of Fatal-Plus® (sodium pentobarbital, Patterson Veterinary Supply, Inc., Devens, MA, USA). Animals were checked for any reflexes to ensure complete unconsciousness before an incision was made to expose the heart. A round needle connected to a cartridge pump, model 7519-06 (Masterflex L/S – Cole-Parmer, Illinois, USA) was inserted and clamped into the left ventricle and the right atrium was then cut to allow fluid drainage. The animal was perfused quickly with cold 0.1 M PBS to clear blood before fixing the brain with cold 4% paraformaldehyde (PFA, Electron Microscopy Sciences, Hatfield, USA). Brains were post fixed in 4% PFA solution overnight (12-24 hr) at 4°C followed by 30% sucrose solution at 4°C overnight for cryoprotection. 40 µm slices were cut on a LEICA SM200R microtome, (Physitemp Instruments Inc., NJ, U.S.A). Brain slices were collected and stored in 0.02% sodium azide in PBS (0.1 M) and mounted on Superfrost Plus Microscope Slides (Fisherbrand – Fisher Scientific, Pittsburgh, U.S.A), coverslipped, and examined using an Olympus Fluoview FV10i Confocal Microscope (Olympus, Shinjuku, Tokyo, Japan).

Tissue from animals that had received viruses carrying shRNAs were examined for expression of GFP and ds-Red to confirm proper placement, infection and recombination of the conditional shRNA constructs. Fiber placement was confirmed by observation of GFP expression and position of the fiber track.

### Fiber Photometry Data Processing

Signal processing and data analyses were carried out in MATLAB and graphed in either MATLAB or GraphPad Prism 9. Raw recorded signals from each assay were converted to change in fluorescence (ΔF) over the average fluorescence for the entire recording (F0). The fluorescence intensity for each point was therefore expressed as a change from average intensity over the complete test recording and expressed as ΔF/F0. To obtain the final ΔF/F0 values corrected for background fluorescence, ΔF/F0 was calculated for both the calcium dependent signal and the control signal and then subtracted to give a corrected ΔF/F0, with F as the signal value of a given sample at any time point and F0 the average of all signal values throughout the whole recording:

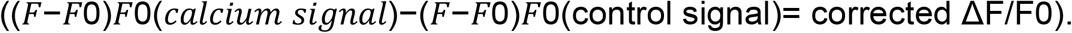

Baseline fluorescence can vary between animals due to difference in sensor expression and/or sensitivity of the fiber. Values for each animal were therefore normalized by z-scoring as follows ^30^:

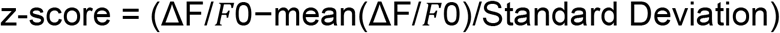

Z-scored ΔF/F0 for each test was time locked with scored behavioral epochs, such as crosses, time in dark compartment and states of immobility.

In the social defeat paradigm, changes in fluorescence were analyzed throughout the pre-attack and social defeat periods. GCaMP fluorescence peak counts during social defeat encounters were counted as the number of peaks that were more than 2 standard deviations from baseline.

For physostigmine injection studies, trendlines were derived for each recording to provide a clearer characterization of the activity pattern using MATLAB with a trending factor of 15000. Error bars were calculated as standard errors of the mean to show variability for each measurement. To ensure visibility on graphs, error bars were not applied to every signal but were applied with a binning factor 8 of the smoothened data.

### Statistical analyses

Statistical analyses were carried out in GraphPad Prism 9 or SPSS17. Results are shown as the mean ± standard error of the mean (SEM). A two-way Analysis of Variance (ANOVA) was used to compare the different targeted neuronal subtypes in the FP paradigm. Two-way ANOVA was also used to analyze behavioral outcomes in nAChR knockdown studies, with “Cre-Line” as a within subject factor, and “knockdown” as a between-subject factor. One-way ANOVA was used to compare locomotor activity of the different Cre-lines in the FP paradigm. Fisher’s Least Significant Difference (LSD) multiple comparisons tests were used for further investigation when a main effect was discovered in the ANOVA.

## RESULTS

### Increased activity of BLA neuronal subtypes during active coping in response to stressors

Activity of excitatory projection neurons in the BLA is thought to be engaged in stressful situations, whereas networks of inhibitory interneurons are thought to be differentially engaged to shape that output. To evaluate the engagement of different BLA neuronal subtypes in behaving animals, fiber photometry of GCaMP6f fluorescence was used to measure the activity of CaMKII-, PV-, SOM- and VIP-expressing neurons throughout all phases of the light-dark and the tail suspension tests, as well as during epochs of social defeat and subsequent social interaction. As might be predicted, in the light-dark box (**Fig. 1C)** overall BLA neuronal activity was significantly higher when animals explored the stressful, bright compartment compared to the dark compartment (**Fig. 1D**; ANOVA: F (1, 30) = 18.41, P=0.0002). When focusing on individual cell types, the same pattern was observed in CAMKII (t(5) = 3.49, P = 0.001), PV (t(5) = 2.13, P = 0.04), and SOM (t(5) = 3.1, P = 0.004) neurons (Fig. 1B). The activity of VIP neurons, however, did not differ when animals were exploring either of the two compartments (t(5) = 0.09, P = 0.92). No significant differences in overall locomotor activity were observed across Cre-expressing mouse lines (**Supp Fig. 1**).

Similar to BLA neuronal activity, ACh levels also increased above baseline when animals explored the light compartment and dipped below baseline when they were in the dark compartment (t(5) = 3.46, P = 0.018; **Fig. 1E**), demonstrating that exploration of a stressful environment increases ACh release in the BLA.

In the tail suspension test (**Fig. 1F**), activity of BLA neurons increased during active (mobility) compared to passive (immobility) coping (**Fig. 1G**; immobility; ANOVA: F (1, 30) = 79.37, P < 0.0001), with no overall interaction by cell type. Indeed, *post hoc* comparisons revealed that CAMKII neurons (t(5) = 4.313, p < 0.001) and all three GABAergic neuronal subtypes (PV: t(5) = 4.2, p > 0.001; SOM: t(5) = 4.6, p > 0.001; VIP: t(5) = 4.7, p > 0.001) exhibited a significant increase in activity during active coping compared to passive coping. Interestingly, activity of all BLA neuronal subtypes began to change before the transition from either active to passive, or passive to active coping (**Fig. 1H and Supp Fig.2**), suggesting that activity in the BLA neuronal network may initiate the change in coping strategy in response to stressful events. Further, because changes in fluorescence preceded changes in movement, this also suggests that movement artifacts are not likely to account for the changes in fluorescence measured in the TST. Similar to BLA neuronal activity, ACh levels increased above baseline during active coping (mobility) and fell below baseline levels during passive coping (immobility) (t(5) = 2.63, P = 0.046; **Fig. 1I**).

During the social interaction test (**Fig. 1J**, there was greater overall BLA neuronal activity when the animal voluntarily moved into the zone with the CD1 mouse, compared to when it was not in proximity to the aggressor (**Fig. 1K**; F (1, 30) = 18.71; p = 0.0002), although there was no interaction with cell type (F < 1). *Post hoc* analyses revealed that only activity of PV and VIP neurons increased significantly (Fig. 2B; PV: t(5) = 2.64, p = 0.01; VIP: t(5) = 3.48, p = 0.002). However, we did not observe a significant change in peaks of neuronal activity between the first and last defeat session, or when activity was compared before and after each defeat encounter (**supp Fig. 3 A, B**; all F < 1), suggesting that the stressful social exposure alone was not sufficient to engage the BLA network and that multiple defeat sessions did not alter BLA activity in response to exposure to the aggressor mouse.

**Figure 2:**
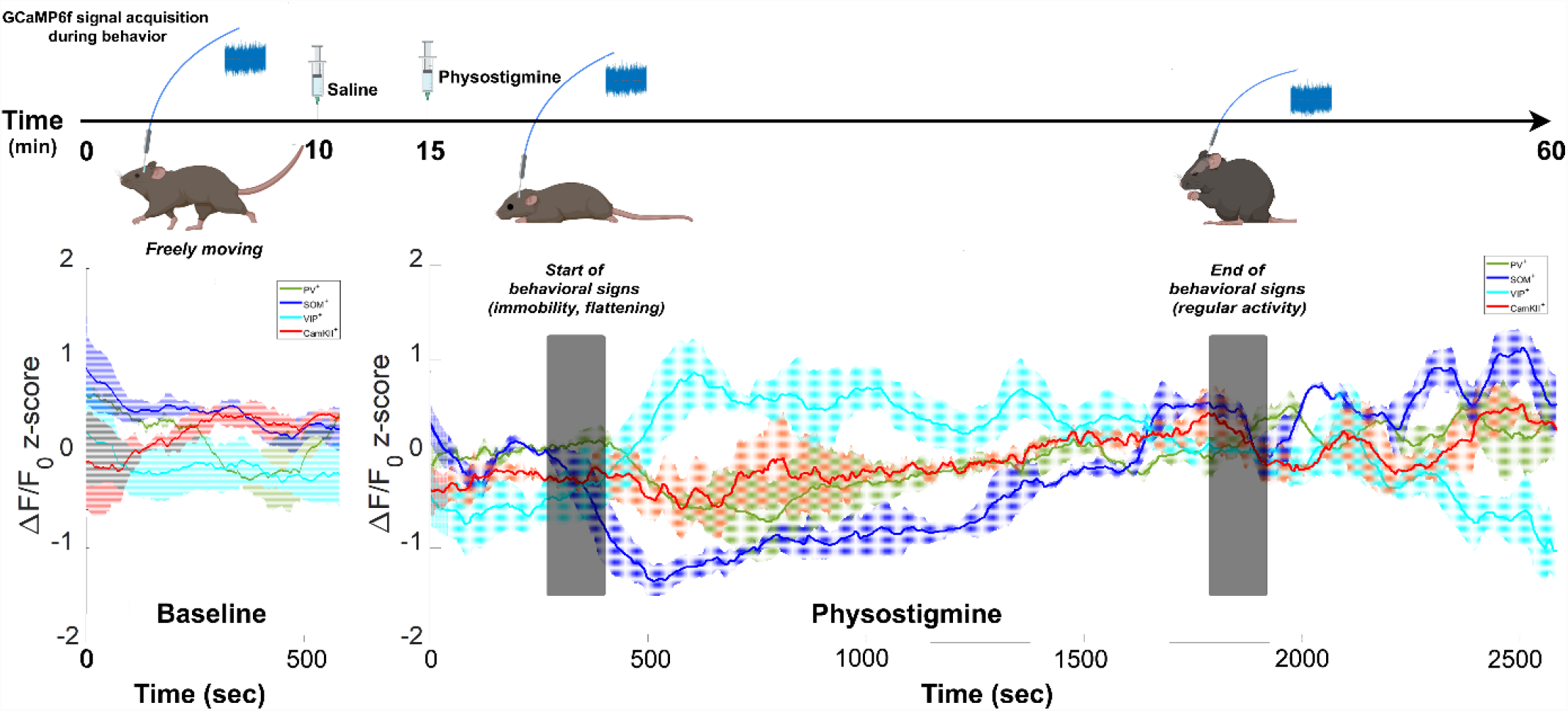
GCaMP6f activity in CaMKII, VIP, PV and SOM neurons following physostigmine i.p. injection (0.15 mg/kg). (**A**) Timeline of injections and measurements in the physostigmine experiment. (**B**) Recording before (Baseline) and after physostigmine injection expressed as z-scored ΔF/F0. Solid lines represent means across animals. Shaded areas represent SEM. N = 5 per group. Partially created with Created with BioRender.com.

When looking at GRAB 3.0 activity, there was high variability, and changes in ACh levels when animals were in the interaction zone did not reach significance (**Fig. 1L**).

### Prolonging ACh signaling alters E/I balance in the BLA network by increasing activity of PV, and decreasing activity of SOM, GABA neurons

In order to determine whether ACh signaling engages activity of the BLA network similarly to stressful stimuli, we evaluated the activity of CaMKII, PV, SOM and VIP neurons in freely-moving mice using fiber photometry to measure GCaMP6f fluorescence (**Fig. 2**) following injection of the cholinesterase inhibitor physostigmine.

Fluorescence measurements reached a stable baseline in all four cell types within ~2 min. Injection of saline resulted in minimal perturbation of activity (**Supp Fig. 4**). In contrast, ~8-10 min after physostigmine injection, when behavioral signs began to be observed (decreased mobility), activity of SOM neurons decreased markedly, before slowly recovering to baseline over ~25 min, when behavioral signs resolved. Activity of SOM-expressing cells continued to increase until the end of testing (60 min after injection). Activity of VIP neurons mirrored the activity of SOM cells, increasing rapidly within 2 min of behavioral signs of physostigmine effects, before slowly returning to baseline ~25 min post-injection once behavioral signs resolved, and continuing to decrease below baseline until the end of observations at 1 hr. In contrast, activity of CaMKII and PV neurons changed only modestly following physostigmine injection. Thus, prolonging ACh signaling changes the balance in activity in the BLA, increasing firing of VIP relative to CaMKII neurons, while decreasing the ratio of SOM/CaMKII neuronal activity.

### Knockdown of nAChR subtypes in GABA neurons, but not principal neurons, increases active coping behaviors in response to stressors

Since ACh release is increased in the BLA in response to stress ^7, 9, 31^, and prolonging ACh signaling with physostigmine alters the balance of activity between CaMKII, SOM, PV and VIP neurons, we next determined whether decreasing ACh signaling in the network by knocking down expression of its receptors in each neuronal subtype would alter stress coping behaviors. We first determined whether changing the overall E/I balance in ACh signaling through nAChRs in the BLA would alter stress relevant behaviors. AAV2 carrying β2 or α7 nAChR shRNAs were therefore injected into the BLA of either CAMKII-Cre or GAD65-Cre mice (**Fig. 3A and B**), and behavioral responses were measured in several stress paradigms. In the light-dark box, there was an overall interaction in time spent in the light compartment between mouse line and knockdown (**Fig. 3C**; F(2, 65) = 5.83, p = 0.005). Further analysis indicated that only GAD-Cre animals showed a significant effect of nAChR knockdown (F(2, 36) = 5.87, p = 0.009) which was driven by a significant effect induced by α7 nAChR shRNA in the BLA of GAD-Cre animals (p = 0.0017). Thus, decreasing α7 nAChR expression in BLA increased active stress coping in the light-dark box. Similarly, in the tail suspension test a main effect was only observed for nAChR knockdown in GAD-Cre mice (**Fig. 3D**; F(2, 38) = 6.75, p = 0.003), and post hoc analysis revealed that decreasing α7 nAChR expression in the BLA of GAD-Cre mice decreased the time immobile, or passive stress coping, in the TST (p = 0.003). In contrast, no effect of perturbing E/I balance of nAChR signaling in BLA was observed in the forced swim (**Fig. 3E**) or the social defeat tests (**Fig. 3F;** all F < 1).

**Figure 3:**
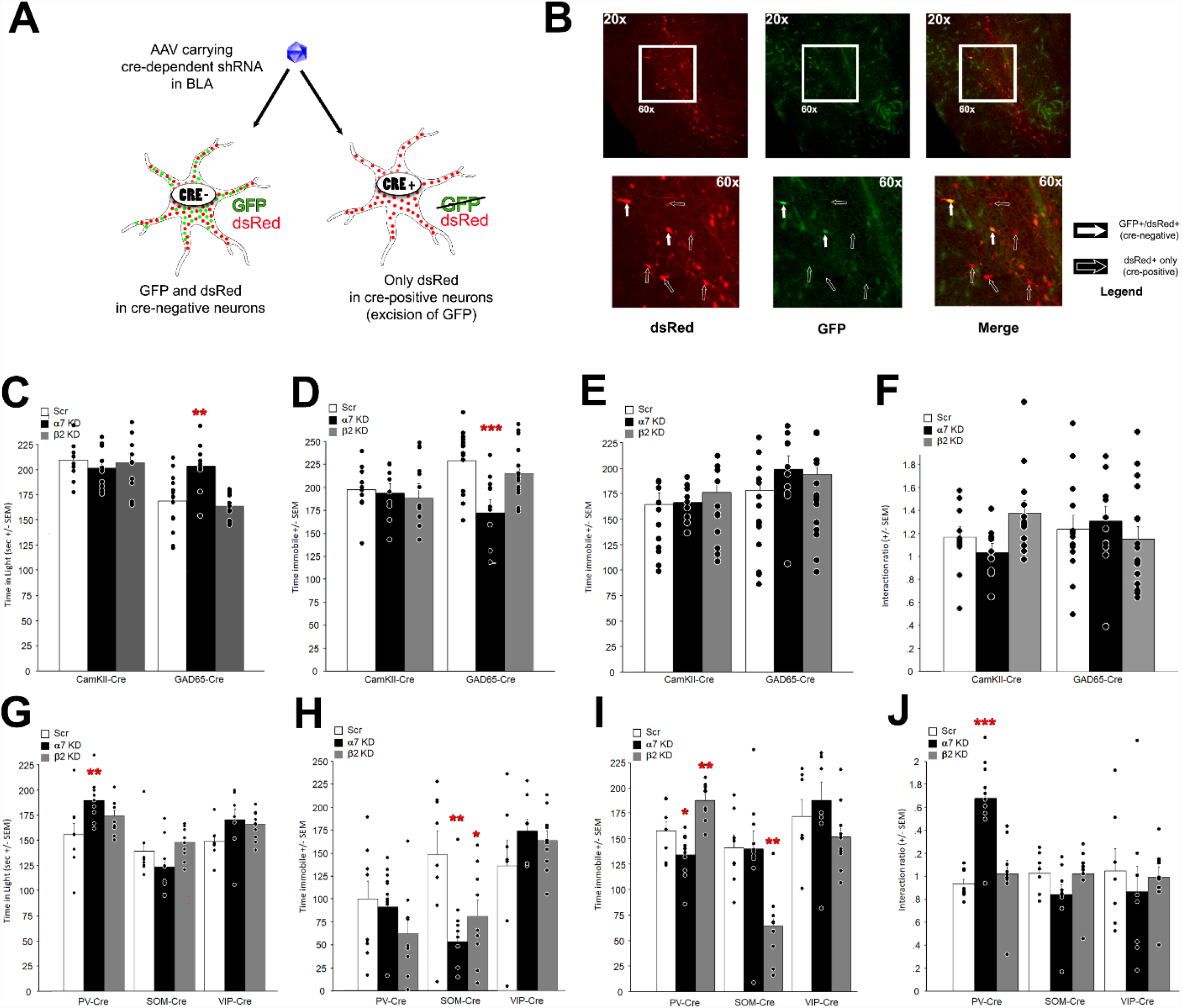
α7 or β2 nAChR knockdown in principal (CAMKII-Cre) or inhibitory (GAD65-Cre) neurons or GABAergic subtypes (VIP-Cre, PV-Cre, SOM-Cre). (**A**). Expression and targeting were checked in all animals by immunohistology following completion of testing. (**B**) Photomicrographs of BLA sections following Cre-dependent knockdown. dsRed and GFP expression demonstrate overall infection. Upon Cre-mediated recombination, GFP is excised and neurons only express dsRed. Cells with both dsRed and GFP expression (solid arrows) were infected but did not express the conditional shRNA (Cre-negative neurons). Cells that only expressed dsRed had excised GFP (hollow arrows) via Cre-mediated recombination and expressed the shRNA (Cre-positive neurons). Effects of α7 or β2 nAChR knockdown in CaMKII or GAD65 neurons in the light-dark box (**C**), tail suspension (**D**), forced swim (**E**), and social interaction after social defeat (**F**). Effects of α7 or β2 nAChR knockdown in PV, SOM or VIP neurons in the light-dark box (**G**), tail suspension (**H**), forced swim (**I**), and social interaction after social defeat (**J**). Bars represent averages across animals; dots represent data from each mouse tested; error bars represent SEM. N = 9-15. * p < 0.05; ** p < 0.01; *** p < 0.001. Partially created with Created with BioRender.com.

### Targeted knockdown of α7 or β2* nAChRs in BLA GABA neuron subtypes confers different patterns of stress resilience across distinct behavioral paradigms

Since perturbing E/I balance of BLA ACh signaling by knocking down nAChRs in GABA neurons, but not principal neurons, increases stress coping, and prolonging ACh signaling with physostigmine has differential effects on different GABA neuron subtypes, we next determined whether knocking down α7 and β2 nAChRs in VIP, SOM, or PV-expressing GABA neurons might have differential effects on stress coping.

In the light-dark box, there was an overall interaction for the time spent in the light compartment between Cre-expressing mouse lines and shRNA infusion (**Fig. 3G**; F(4, 71) = 3.36, p = 0.015). *Post hoc* analyses revealed that knocking down α7 nAChR in PV neurons increased the time spent in the light compartment (p = 0.006). Thus, knockdown of α7 nAChR in PV neurons confers active stress coping in the light-dark box and recapitulates what was seen for knockdown of α7 nAChR in GAD65 neurons (**Fig. 3C**).

In the tail suspension test, a similar overall interaction was observed, albeit with more variability (**Fig. 3H**; F(4, 69) = 4.02, p = 0.054). *Post hoc* analyses indicated that knockdown of α7 or β2* nAChRs in SOM-expressing neurons significantly reduced the time spent immobile (p = 0.001 and p = 0.02, respectively).

A similar statistical interaction was observed for the time spent immobile in the forced swim test between mouse lines and knockdown (**Fig. 3I**; F(4, 69) = 8.53, p = 0.001), with significant differences only detected in PV and SOM neurons. Indeed, α7 nAChR knockdown in PV neurons decreased the time spent immobile (p = 0.03) as did β2 nAChR KD in SOM neurons (p = 0.002). Surprisingly, the β2 nAChR shRNA significantly *increased* the time spent immobile in the forced swim test (p = 0.01), which was unexpected. Finally, following social defeat, there was a Cre-line by shRNA interaction in the social interaction test (**Fig. 3J**; F(4, 70) = 6.52, p = 0.0002) but *post hoc* analyses indicated that only α7 knockdown in PV neurons increased the interaction ratio (p = 0.0001), suggesting increased active stress coping following social defeat. nAChR knockdown in VIP neurons did not significantly change behavior in any of the tests.

## DISCUSSION

In the current study we measured activity of excitatory and inhibitory neuronal subtypes in the BLA of animals subjected to acute and chronic stressors using GCaMP6f via fiber photometry. We found that for the most part, activity of both excitatory and inhibitory neuronal subtypes increased during stressful challenges, and that this activity was most apparent during active stress coping. Because acetylcholine signaling is a critical modulator of stress reactivity ^10, 32, 33^, we also measured ACh levels using a GRAB_ACH_ sensor and found similar increases during active coping behaviors. We then investigated the activity of neuronal subtypes in the BLA network following the administration of physostigmine, an anticholinesterase inhibitor that can induce symptoms of anxiety and depression in humans and in animal models ^28, 34, 35^ and found that prolonging ACh signaling perturbed E/I balance by altering signaling between specific inhibitory neuronal subtypes, increasing activity of VIP neurons and decreasing the activity of SOM neurons relative to CaMKII neurons. Since stress increases ACh signaling ^9, 31^, and because decreasing nAChR signaling within the amygdala increases stress resilience and increases stress coping behaviors ^10^, we hypothesized that knockdown of nAChRs in specific BLA neuronal subtypes might restore E/I balance during stress exposure and result in increased active stress coping. Interestingly, only nAChR knockdown in inhibitory neurons increased active stress coping, although different neuronal and nAChR subtypes contributed across stressful paradigms.

GABAergic interneurons provide critical regulation of excitatory output that is essential for the processing of information throughout the brain ^36^. Across the behavioral paradigms used here, the broad pattern of activation of neuronal subtypes and release of ACh in BLA was similar, with greater activity during challenging situations (light side of the light dark box and during social interaction after social defeat) and during active coping (mobility during TST and FST). Of note, some test-specific patterns were also observed. For instance, VIP neuron activity remained unchanged in the bright side of the dark box. While we cannot exclude an experimental false negative, this may suggest that the “challenge” of being in a bright environment is not sufficient to engage BLA VIP neurons, which would also limit inhibitory inputs from VIP neurons onto CAMKII, SOM and PV neurons, overall disinhibiting the network ^37, 38^. A fiber photometry study evaluating neuronal activity in the prefrontal cortex (PFC) showed that SOM and PV neurons were more active when animals explored the open arms of an elevated plus maze ^39^ similar to what we observe in the BLA in the current study. However, this study also found that VIP neurons were more active in the open arms and that inhibiting PFC VIP neurons increased time spent in the open arms. It is likely that GABAergic networks show differential patterns of activity across brain regions ^40^, however, VIP neuron activity could increase depending on the severity of the stressful stimulus. This possibility is in line with data in the current study showing that VIP neuronal activity increases significantly in the TST, a test that involves an inescapable stressor, in contrast to voluntary exploration in the light-dark box ^41–43^. In this paradigm, we found that all BLA neuronal subtypes measured, including VIP neurons, showed higher calcium signaling during active coping, when the mice were struggling to escape. Conversely, neuronal activity of all BLA cell types was below baseline when animals were immobile. Strikingly, changes in activity of all BLA neuronal subtypes *preceded* the transitions between behavioral states (from immobile to mobile and vice versa), suggesting that activity of the BLA network could drive active coping behavior in response to stress, rather than solely being a marker of stress severity.

Release of ACh in BLA also occurred in response to stressful environmental stimuli, but only when the situation involved active coping. This is interesting because it suggests that under conditions in which stress is controllable (light-dark exploration, social interaction after defeat), ACh release does not drive maladaptive behavior but in fact may contribute to resilience to stressful challenges. This is consistent with the idea that optimal ACh release increases signal-to-noise ratios, heightens threat perception, and favors adaptative coping, whereas prolonged ACh release leads to maladaptive attention to negative stimuli and biased encoding of stressful experiences ^44^.

In this study we did not observe any changes in BLA neuronal activity, as measured by counting peaks of calcium fluorescence before and after social defeat, and between the 1^st^ (including before the first defeat) and the 8^th^ defeats. This suggests that repeated social defeat exposure did not sensitize the BLA network to the proximity of an aggressor. Since recording was not done during physical attacks (to avoid potential damage to the fiber), it is possible that changes in BLA neuronal activity may only be observed when an active threat is present, although proximity of an aggressor is thought to be a significant stressor. In contrast, during the social interaction test at the end of the defeat paradigm, neuronal activity increased, at least modestly, in all BLA cell types measured when the animal was in the interaction zone with an aggressor mouse, although the increase in activity was only significant for PV and VIP neurons. As in the light-dark test, the volitional approach to the stressor (in this case the CD1 mouse) might lessen its intensity compared to the inescapable stress of the TST, which may also be reflected in the non-significant fluctuation of ACh levels during social interaction.

In contrast to naturalistic stressors, prolonging ACh signaling with physostigmine increased activity of BLA VIP neurons selectively, while decreasing activity of SOM neurons. The observation that VIP neurons are a primary target for prolonged ACh signaling in the BLA is consistent with optogenetic stimulation of ACh terminals in BLA slices showing that fast nAChR currents occur in late-firing interneurons, but not fast-spiking interneurons, whereas principal neurons largely receive muscarinic receptor innervation that alters firing in a state-dependent manner depending on firing rate ^45^. The decrease in SOM neuron firing following physostigmine administration could be a result of ACh-mediated VIP neuron firing. Studies in BLA ^46^ and neocortex ^38, 47^ show that VIP interneurons are activated by aversive stimuli ^19, 46^ and inhibit both PV and SOM interneurons, in turn leading to disinhibition of projection neurons. Following physostigmine administration, activity of PV and CaMKII neurons in BLA increased modestly, until behavioral signs were no longer observed. Thus, prolonged ACh signaling shifts E/I balance in the BLA, in contrast to the relatively consistent increase in activity of both excitatory and inhibitory BLA cell types in response to environmental stressors. This suggests that perturbations that result in prolonged changes in ACh signaling could bias responses to stress. It should be noted that a caveat of this experiment is that systemic administration of physostigmine prolongs ACh signaling throughout the brain and body. It is therefore possible that the changes in neuronal activity observed in the amygdala may be due to alterations in activity of other neuronal systems ^48, 49^ and not to direct effects of ACh in the BLA.

In order to determine which nAChR subtypes contribute to the ability of prolonged cholinergic stimulation to alter behaviors dependent on E/I balance in the BLA, we knocked down expression of α7 or β2* nAChRs in principal neurons (CaMKII) and GABA (GAD-65+) neurons as a group, and then in each of the GABAergic subtypes measured in fiber photometry studies (SOM, PV, VIP). Decreasing activity or expression of α7 or β2 nAChRs in a non-cell type-selective manner in the BLA can increase resilience and active stress coping in mice ^10^. Consistent with the modest changes in activity of CaMKII neurons in response to physostigmine, knockdown of α7 or β2 nAChRs in principal neurons did not result in significant changes in stress coping in the behavioral paradigms tested. While this was at first somewhat surprising, since excitability of BLA principal neurons is generally thought to be critical for stress sensitivity ^50^, it is also consistent with the finding that principal neurons show relatively modest nAChR responses and are predominantly regulated by muscarinic receptors ^45^.

In contrast, downregulating α7 nAChR expression in GABAergic neurons increased time spent in the light side of the LD box and decreased time spent immobile in the TST but had no effect on behavior in the forced swim or the social interaction tests. These paradigm-specific results are similar to the effects of global α7 nAChR KD in the BLA ^10^, suggesting that effects of α7 in BLA on stress-relevant behaviors are largely driven by effects of ACh on GABA neurons. In contrast, knockdown of β2* nAChRs in CaMKII or GAD-65 neurons had no significant effects. It is possible that β2* nAChRs regulate activity of both principal neurons and inhibitory neurons, and that global knockdown was more effective in altering network activity. Along the same lines, the reciprocal inhibition between different GABAergic cell types ^46^ could result in paradoxical effects of β2* nAChR knockdown on the network. Thus, we further investigated whether knocking down nAChRs in specific GABAergic subtypes might have differential effects depending on cell type, as has been observed in fear learning ^46, 51^.

Effects of nAChR knockdown were cell-type, receptor-type, and test specific, suggesting that nAChR-mediated modulation of the BLA inhibitory network occurs at multiple levels and is not easily decoded *in vivo*. Among the behavioral effects we observed were a significant increase in time spent in the light side of the light-dark box and an increase in social interaction following social defeat following α7 KD in PV neurons, decreased immobility in the tail suspension test following α7 KD in SOM neurons, and decreased time spent immobile in the tail suspension and forced swim test following β2 KD in SOM neurons. Paradoxically, β2 KD in PV neurons resulted in greater immobility in the forced swim test, which may reflect a role for this nAChR subtype in PV neuron activity that is normally balanced by β2* nAChRs on other BLA neuronal subtypes. Perhaps the opposing effects of β2* nAChR KD in SOM and PV neurons on immobility in the forced swim test explains the overall lack of effect of β2* nAChR KD in all GAD65-expressing neurons. Surprisingly, selective nAChR KD in VIP neurons had no behavioral effects across paradigms tested, despite the increased firing of these neurons following physostigmine administration. This could suggest that muscarinic receptors are more important for regulation of VIP neuron firing, or that nAChR signaling on other BLA neuronal subtypes contributes more significantly to behavioral outcomes. For example, PV and SOM neurons are downstream of VIP neurons in the BLA network, so nAChRs on these inhibitory neuronal subtypes may predominate and be more effective in modulating stress-relevant behaviors. One caveat for these studies is that the different Cre-recombinase expressing mouse lines used exhibit somewhat different behavioral baselines (for instance, low baseline immobility in PV-Cre mice was observed in the tail suspension test) which could obscure the consequences of nAChR knockdown. However, taken together cell type-selective nAChR knockdown studies indicate that nAChR-mediated modulation of signaling in the GABAergic network of the BLA does contribute to regulation of E/I balance and is important for behavioral responses to stress. Consistent with a modulatory role, multiple nAChR subtypes have effects on GABA neuron signaling, and effects of nAChRs on distinct GABAergic interneuron subtypes contribute to stress resilience and coping, depending on the type of stressor.

In conclusion (**Fig. 4**), the fiber photometry studies presented here suggest that ACh is transiently released in the BLA and principal and local inhibitory neurons of the BLA increase their activity in response stressful challenges, maintaining E/I balance in the structure in response to physiological stressors. The change in BLA neuronal activity in advance of behavioral transitions suggests that BLA activity may drive active stress coping strategies. In contrast, prolonged ACh signaling following blockade of acetylcholinesterase by physostigmine increases BLA VIP signaling and alters E/I balance. nAChR signaling appears to be more effective in GABAergic neurons of the BLA; however, regulation of the network is complex, and appropriate E/I balance may require nAChR expression on both principal neurons and GABA neurons. Since physostigmine had the greatest effect on BLA VIP neuron firing but nAChR knockdown in BLA VIP neurons had no behavioral consequences in tests of stress coping, muscarinic signaling may be more important for VIP neuron firing, whereas nAChR signaling is more critical in SOM and PV neurons. Taken together, these results provide new insights into the role of ACh in regulation of the BLA GABAergic network in response to stressful stimuli, identifying potential molecular targets for interventions to increase stress resilience and coping.

**Figure 4:**
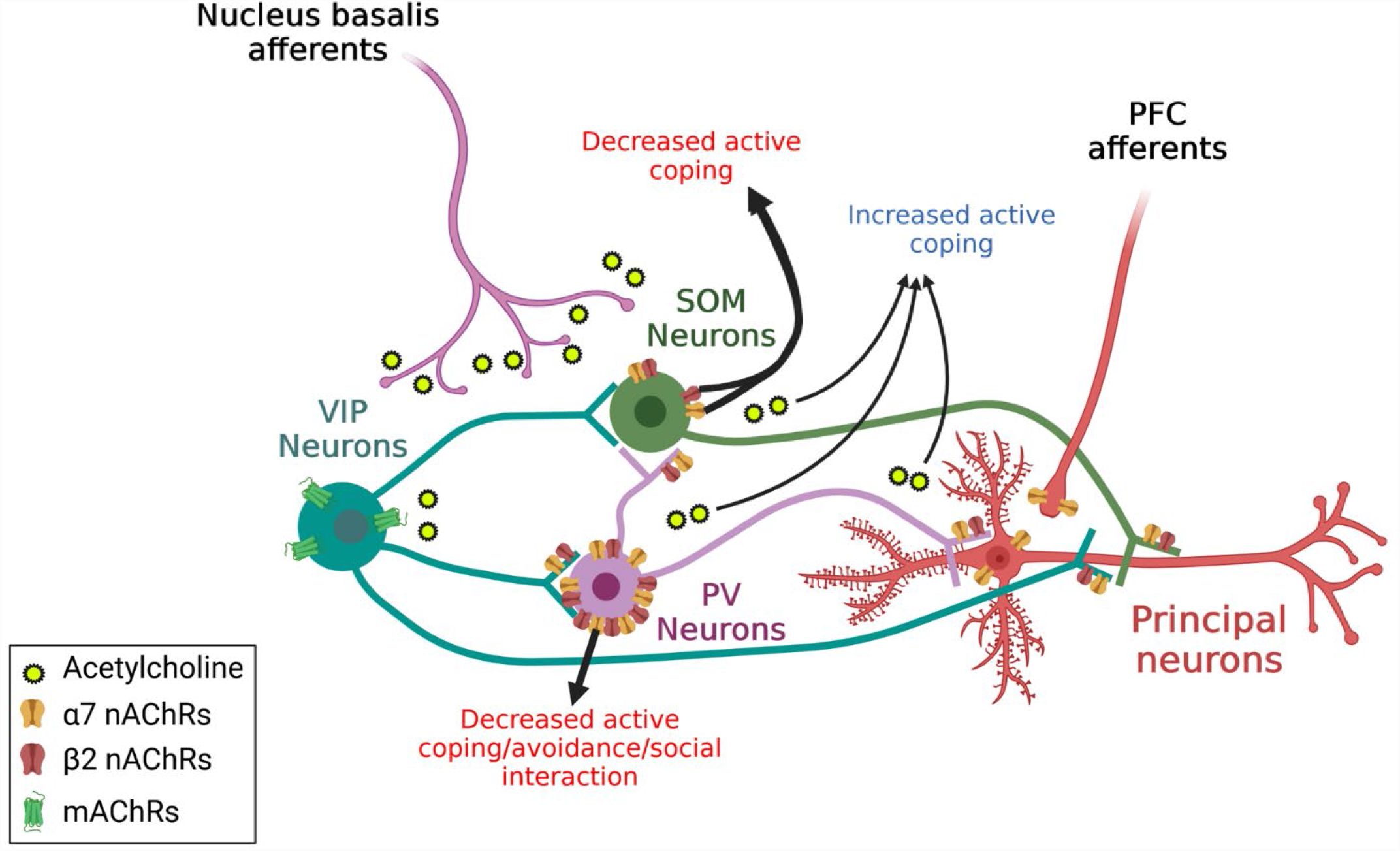
Schematic summarizing effects of ACh and cholinergic receptors in the BLA and their influence on behavior. During passive coping, little ACh is released from cholinergic afferents. Upon transient release of ACh and other neurotransmitters following moderate or transient exposure to stress, GABAergic interneurons and principal neurons increase their activity in concert, enhancing signal-to-noise in the network and increasing coping behaviors. Following prolonged or dysregulated ACh release, E/I balance is disrupted in the network, leading to asynchrony and maladaptive behavioral responses. Created with Biorender.com

## ACKNOWLEDGMENTS

These studies were supported by National Institutes of Health grants MH077681, MH105824 and DA033945 from the National Institutes of Health and a NARSAD Distinguished Investigator grant from the Brain and Behavior Research Foundation. This work was funded in part by the State of Connecticut, Department of Mental Health and Addiction Services, but this publication does not express the views of the Department of Mental Health and Addiction Services or the State of Connecticut.

## Supplementary Figures

**Supp Fig. 1:**
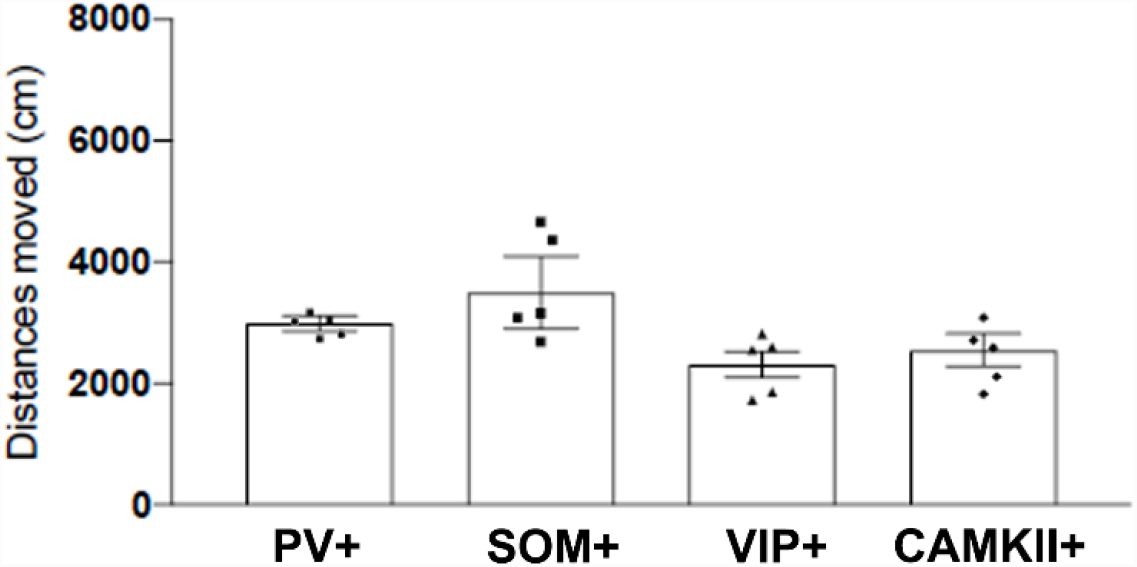
Locomotor activity in an open field of individual Cre-line of mice. Individual dots are average per mouse. Error bars are SEM.

**Supp Fig. 2:**
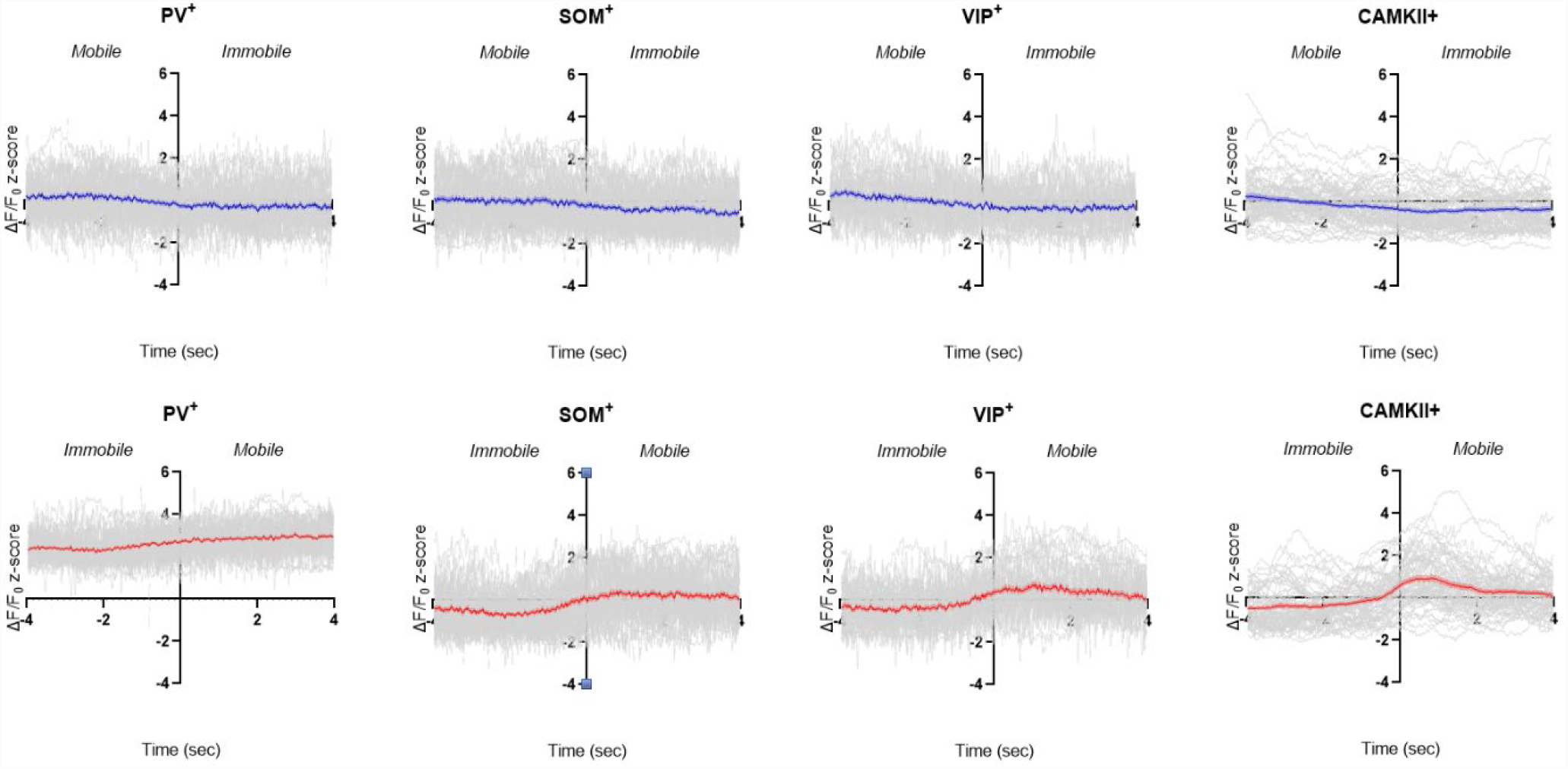
Individual traces of z-scored ΔF/F0 GCaMP6f activity, in each cell type, for each transition (in gray). Colored lines are averages. N = 5 per group.

**Supp Fig. 3:**
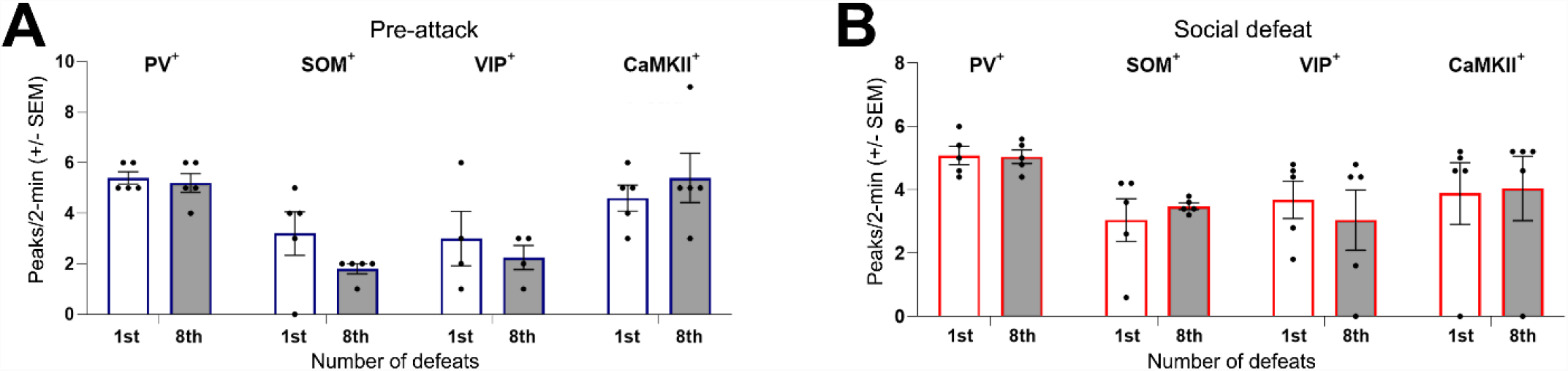
Average peaks counts of z-scored ΔF/F0 GCaMP6f activity, in each cell type, for each transition (in gray), before the attack (**A**) and during social defeat (**B**). N = 5 per group.

**Supp Fig. 4:**
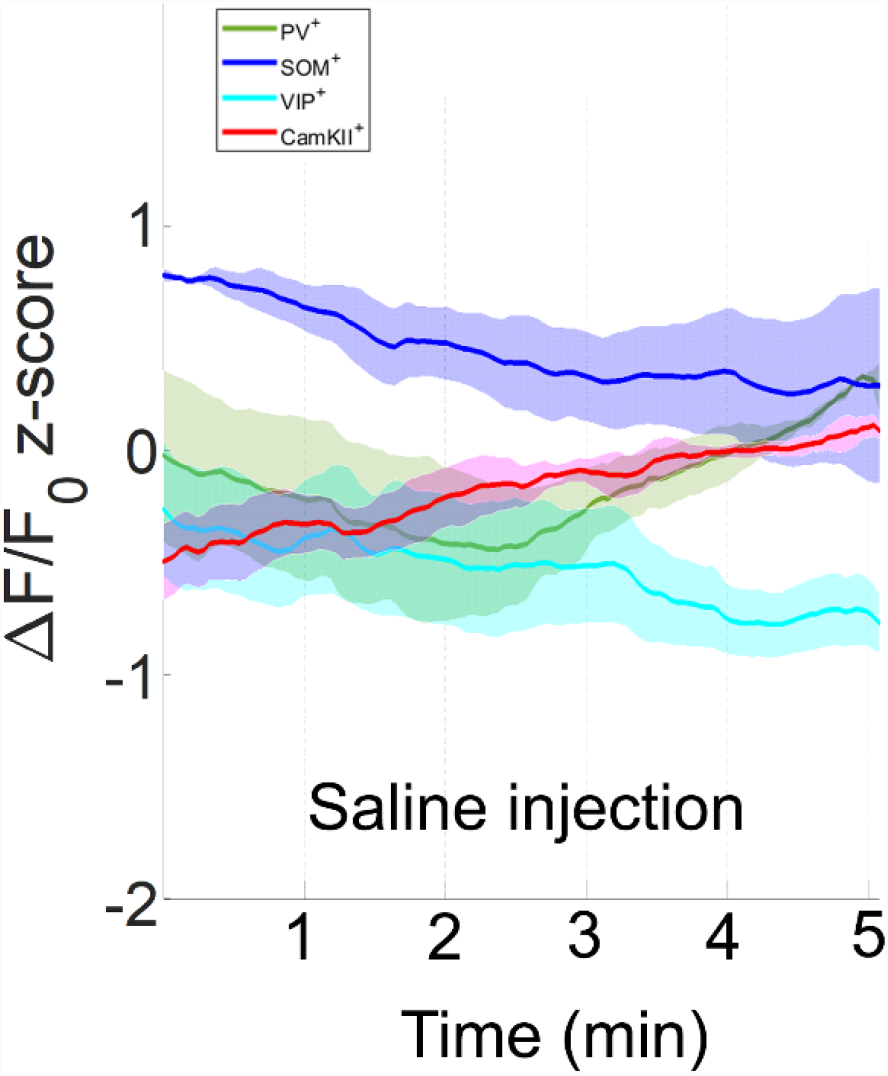
Z-scored ΔF/F0 GCaMP6f activity following Saline injection. Solid lines represent means across animals. Shaded areas are SEM. N = 5 per group.

